# Genome-wide association studies identify new candidate genes and tissues underlying resistance to a natural toxin in drosophilids

**DOI:** 10.64898/2025.12.01.691102

**Authors:** Michele Marconcini, Caroline Fragnière, Ambra Masuzzo, Richard Benton

## Abstract

Many insects can rapidly evolve resistance to artificial insecticides through changes in toxin target proteins. Over longer timescales, insects have also evolved resistance to naturally-occurring toxins to exploit new ecological niches, but much less is known about the mechanisms underlying such adaptations. A classic example is *Drosophila sechellia*, an extreme specialist for the ripe noni fruit of *Morinda citrifolia*, which is toxic for other insects – including the close relatives *D. simulans* and *D. melanogaster* – due to noni’s high content of octanoic acid (OA). The mechanistic bases underlying susceptibility and resistance to OA of different species remain unclear. Here, we first show that the species-specific tolerance of OA is independent of these drosophilids’ distinct microbiomes, reinforcing the notion that this trait is genetically encoded. Screening large, genetically-diverse panels of *D. melanogaster* and *D. simulans* strains revealed broad variation in OA resistance, with some lines surviving as well as *D. sechellia*. Resistance to OA does not correlate with resistance of these lines to other insecticides, implying a distinct toxicity mode-of-action. Genome-wide association and transcriptome-to-phenotype analyses identified multiple genes linked to OA resistance. These genes have diverse expression patterns and functions, including proteins involved in epithelial septate junction formation, lipid transport and tracheal morphogenesis. Loss-of-function analysis in *D. melanogaster* confirmed that at least two of these – Bez, a CD36-family fatty acid transporter, and CG13003, a putative extracellular matrix component – positively contribute to OA resistance. Integration of our findings with those from previous complementary genetic approaches supports a model in which OA has no singular target, and that resistance to this toxin is defined by multigenic and multi-tissue defense mechanisms.

## Introduction

Animals are exposed to myriad volatile and non-volatile chemicals. Some are exploited as beneficial signals, for example, when locating food (Roura and Foster, 2018; Ruedenauer et al., 2023) or conspecifics (Wyatt, 2014). But many chemicals are toxic, such as heavy metals or those produced by plants as defense against herbivores (Erb and Reymond, 2019; Jones et al., 2022). A major line of evasion of intoxication by animals is through chemosensory-mediated avoidance (Wooding et al., 2021; Xiao et al., 2022). However, behavioral responses are often inadequate to escape exposure, and toxic chemicals end up inside animals. Understanding how chemicals enter bodies (e.g., passive absorption, ingestion), the mechanisms underlying their movement between cells, tissues and organs (e.g., diffusion, carrier/transporter proteins), whether they have unique targets or exert general affects (e.g., on cellular membranes), and how animals defend against such toxins (e.g., enhanced body shielding, toxin metabolism/excretion) are important issues spanning the fields of chemical ecology, neuroscience, physiology and evolution.

The susceptibility and resistance mechanisms of insects to toxic chemicals is of particular interest, both because these processes reflect the insect perspective of their arms race with plant food sources (Erb and Reymond, 2019; Jones et al., 2022) or venomous predators (Smith et al., 2013), and because humans have long used chemicals to combat insects that damage our agriculture or act as vectors of pathogens (Umetsu and Shirai, 2020). The best-characterized natural and artificial insecticides are those that have a single, main cellular target (Sparks et al., 2020). For example, cardiac glycosides, produced by plants as defensive compounds by plants, exert their effect through inhibition of the sodium-potassium ATPase pump (Agrawal et al., 2012). Several insect orders have convergently evolved resistance to these toxins through amino acid substitutions in the α-subunit of this pump (Agrawal et al., 2012; Karageorgi et al., 2019). Plant-produced terpenoids are neurotoxic for insects through their antagonism of GABA receptors, and insects have evolved resistance through various mutations in this target receptor. Remarkably, similar mutations have arisen to confer resistance to the artificial insecticide dieldrin, which also targets these receptors (Guo et al., 2023). Indeed, many of the most widely-used synthetic insecticides are agonists or antagonists of neuronal receptors or ion channels (Sparks et al., 2020). While these can be highly effective in killing insects, mutations conferring resistance on the target protein can be rapidly selected in insect populations (Carson, 1962; Sparks et al., 2020). This phenomenon is most famously exemplified by dichlorodiphenyltrichloroethane (DDT), which prevents closure of voltage-gated sodium channels, an effect that has been circumvented in multiple insect orders through “knockdown resistance” (kdr) mutations in the corresponding genes (Busvine, 1951; Dong, 2007).

The mode of action and resistance mechanisms to many toxins remain less clear, typically where chemicals likely have more general effects on insect tissues and resistance cannot be acquired through changes in single genes. Such cases are commonly seen where species have adapted to toxic ecological niches, and provide interesting models to explore broader toxicology mechanisms and evolution of resistance in animals (Despres et al., 2007). For example, plant glucosinolates are potent defense compounds that chemically conjugate to exposed nucleophilic residues in diverse proteins leading to cellular and oxidative stress (Hopkins et al., 2009). The leaf-mining drosophilid *Scaptomyza flava* has evolved tolerance of such compounds, at least in part through duplication and neofunctionalization of glutathione-S-transferase-family detoxification enzymes (Gloss et al., 2014).

One long-recognized example of natural toxin resistance accompanying ecological specialization is the fruit fly *Drosophila sechellia*, which feeds and breeds exclusively on the acid-rich, toxic “noni” fruit of the shrub *Morinda citrifolia* (Auer et al., 2021; Farine et al., 1995; Jones, 2005; Pino et al., 2009). The main noni toxin is thought to be octanoic acid (OA), which is lethal – through an unknown mechanism – to both very closely-related drosophilid species, such as *Drosophila simulans* and *Drosophila melanogaster* (Figure 1A) (Farine et al., 1995; Legal and Plawecki, 1995), and more divergent insects, such as cockroaches, bees and ants (Legal and Plawecki, 1995). *D. sechellia*’s resistance to OA allows it to exploit ripe noni as a unique ecological niche (Amlou et al., 1997; Legal et al., 1994; R’Kha et al., 1991) thereby potentially escaping competition and parasitoidization (Salazar-Jaramillo and Wertheim, 2021).

**Figure 1.**
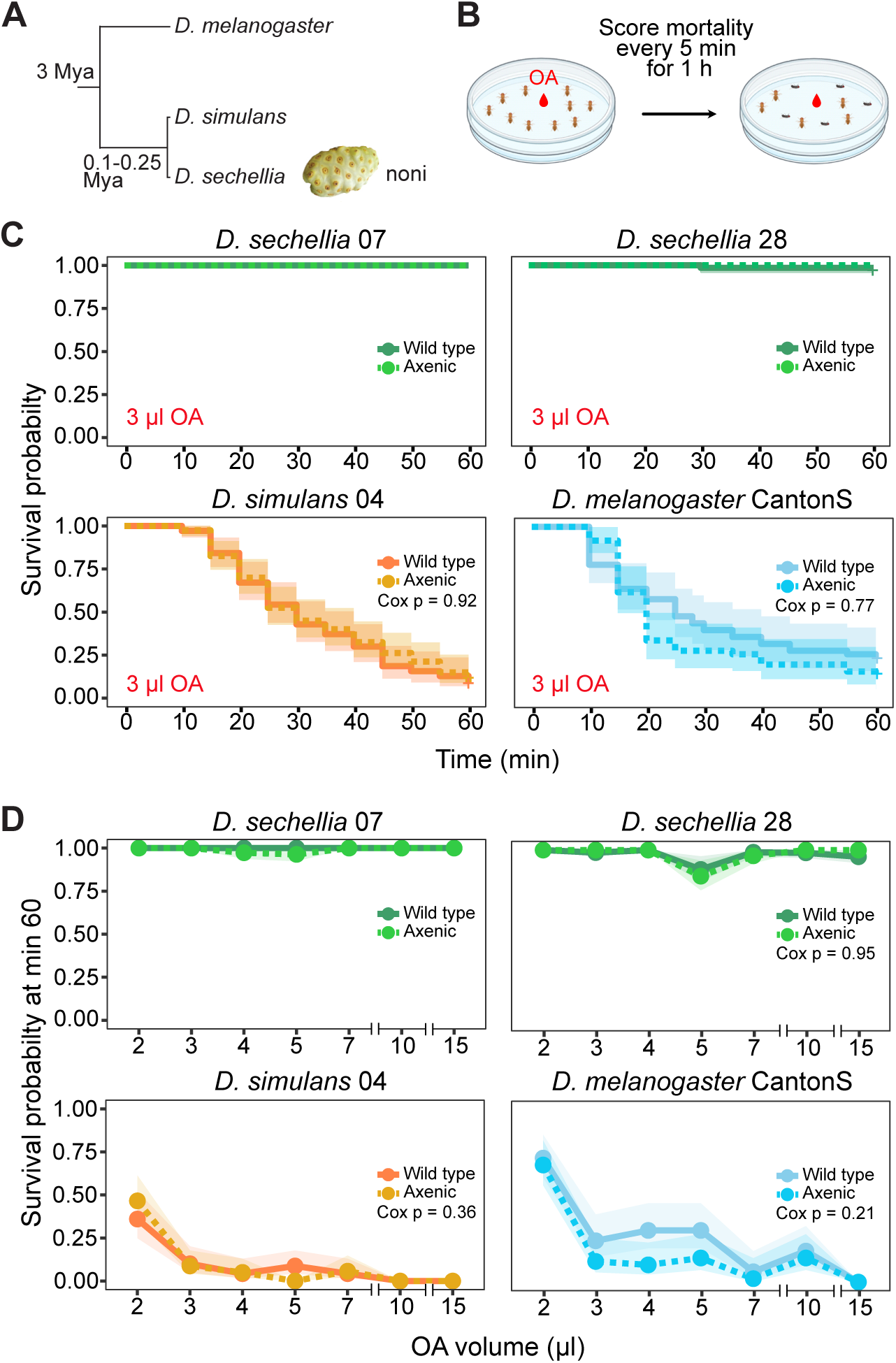
Octanoic acid resistance in axenic flies. (A) Phylogeny of the *Drosophila melanogaster* species complex used in this study. Mya: million years ago. (B) Schematic of the octanoic acid (OA) resistance plate assay. (C) Survival probabilities (y-axis) of axenic adult flies compared to wild-type strains to 3 μl OA over 60 min of exposure (x axis). (D) Survival probabilities (y-axis) of axenic adult flies compared to wild-type strains after 60 min of exposure to different volumes of OA (x-axis). Each replicate for (C,D) consisted of ten 2-7 day-old female flies. N replicates ≥ 5, n flies ≥ 50. Shaded areas represent the 95% confidence intervals. P-values are not reported when no mortality occurred for one or both groups (i.e., wild type and axenic flies). Raw data are available in File S1.

Pioneering attempts to analyze resistance to OA (or noni fruit) used quantitative trait locus (QTL) mapping in hybrids of *D. sechellia* and *D. simulans* at different life stages (Huang and Erezyilmaz, 2015; Jones, 1998, 2001). These efforts showed that at least five genomic loci contribute to resistance, including a major-effect region on chromosome arm 3R and additional loci on all other main chromosome arms (Auer et al., 2021; Huang and Erezyilmaz, 2015; Jones, 1998, 2001). While these regions are too large to pinpoint specific genes, subsequent fine mapping of the 3R QTL identified a ∼170 kb region, containing 18 genes.

Testing of these candidates by RNAi in *D. melanogaster* identified three (*Osi6*, *Osi7*, *Osi8*) as modulators of OA resistance (Andrade Lopez et al., 2017; Hungate et al., 2013; Lanno et al., 2019b), though if and how they contribute to OA resistance in *D. sechellia* awaits further investigation. Complementary studies employed comparative bulk transcriptomics of *D. sechellia* and other drosophilids in control and noni- or OA-exposed conditions (Dworkin and Jones, 2009; Lanno et al., 2017) to identify candidate toxin metabolism genes by virtue of their higher basal expression in *D. sechellia* and/or their induction by OA/noni presentation. Of a large number of candidates, the esterase Est-6 was implicated in OA tolerance through RNAi in *D. melanogaster* (Lanno et al., 2019a). Recently, we used experimental evolution of *D. simulans* and genome-wide CRISPR-based screening approaches in *D. melanogaster* cell lines to identify other genes contributing to OA resistance (Marconcini et al., 2025). Although loss of two of these in *D. sechellia* – the putative detoxification enzyme Kraken and the mitochondrial metabolic regulator Alkbh7 – indicates that they contribute to the natural OA resistance of this species (Marconcini et al., 2025), it is clear that they are only part of the resistance mechanism. In this study we sought to identify additional candidate loci that contribute to OA susceptibility and resistance in drosophilds through complementary, cross-species genetic approaches.

## Results

### OA susceptibility and resistance in drosophilids is independent of the microbiome

The high resistance of *Drosophila sechellia* to OA/noni has been assumed to reflect genetic adaptation of this species (Huang and Erezyilmaz, 2015; Jones, 1998). However, resistance of various insect species to several toxic chemicals has been linked to the microbiome (Brown et al., 2021; Malook et al., 2025), through presumed microbial functions in chemical detoxification or modulation of host gene expression and physiology. Comparative studies of drosophilid microbiota indicate that *D. sechellia* collected from noni fruit has a distinctive microbiome, dominated by a single *Lactobacillales* taxon, but otherwise an extremely low bacterial community richness and evenness (Chandler et al., 2011; Heys et al., 2021). Moreover, when *D. melanogaster* was fed food containing the *D. sechellia* microbiome, it lost its normal aversion to OA (Heys et al., 2021), raising the possibility that the *D. sechellia* microbiota can influence host behavior.

We therefore first investigated whether the distinctive microbial profile of *D. sechellia* contributes to this species’ resistance to OA by generating axenic flies, through culture on a cocktail of antibiotics (see Methods). Given the low diversity of the *D. sechellia* microbiome, and the observations that microbes can sometimes increase toxicity through metabolic “activation” (Peterson, 2024), we also generated axenic *D. simulans* and *D. melanogaster*. These flies, as well as untreated controls, were tested for OA resistance in a “plate assay” (Jones, 1998; Marconcini et al., 2025), in which mortality of flies exposed to a droplet of OA is monitored over the course of 1 h (Figure 1B). As expected, when exposed to 3 μl OA, the two *D. sechellia* strains showed almost no mortality, whereas both *D. simulans* and *D. melanogaster* had very few survivors by the end of the experiment (Figure 1C). Importantly, no differences in survival probability were observed between axenic and control strains for any species (Figure 1C). We then extended the assay to a broader range of OA volumes and found no evidence of differences in survivorship between axenic and control flies (Figure 1D). These results suggest that the microbiome of these drosophilid species does not make a measurable contribution to OA resistance, reinforcing the idea that this is an exclusively, genetically-determined trait.

### Phenotypic variation in OA resistance across genetically-diverse *D. melanogaster* and *D. simulans*

One effective approach to identifying the genetic basis of complex traits is to examine the relationship between genetic and phenotypic variation, as implemented in genome-wide association studies (GWAS). However, GWAS analyses require both substantial standing genetic variation and adequate statistical power, which is largely determined by the number of available genotypes or strains. These requirements are difficult to meet in *Drosophila sechellia*, as no large strain panel exists and the species is over an order of magnitude less polymorphic than other members of the *melanogaster* subgroup (Legrand et al., 2009). Commensurately, OA resistance appears similar across tested laboratory and wild-caught strains (Shahandeh). Study of a small number wild-type *D. melanogaster* and *D. simulans* strains has revealed the existence of intraspecific variation in sensitivity to OA (Figure S1A and (Andrade Lopez et al., 2017; Colson, 2004)). Given the phylogenetic proximity of these species to *D. sechellia* (Figure 1A), we reasoned that we might uncover novel loci contributing to OA resistance by taking advantage of the substantial genetic diversity within large collections of sequenced, isogenic strains of *D. melanogaster* (the *Drosophila melanogaster* Genetic Reference Panel (DGRP) (Mackay et al., 2012), originating from a fruit market in Raleigh, North Carolina) and *D. simulans* (Signor et al., 2018) (originating from the Zuma organic orchard, California).

To analyze OA resistance in these large panels, we used a simpler “tube assay” (Amlou et al., 1997; Colson, 2004; Marconcini et al., 2025), in which ∼25 flies were placed in culture vial containing a tissue paper spotted with 3 µl of OA in 3% glucose solution, and scored viability after 24 h (Figure 2A). In previous studies of various non-*D. sechellia* drosophilids (Amlou et al., 1997; Colson, 2004; Marconcini et al., 2025), such a quantity of OA resulted in lethality of approximately half of the animals, making it an effective dose for screening purposes.

**Figure 2.**
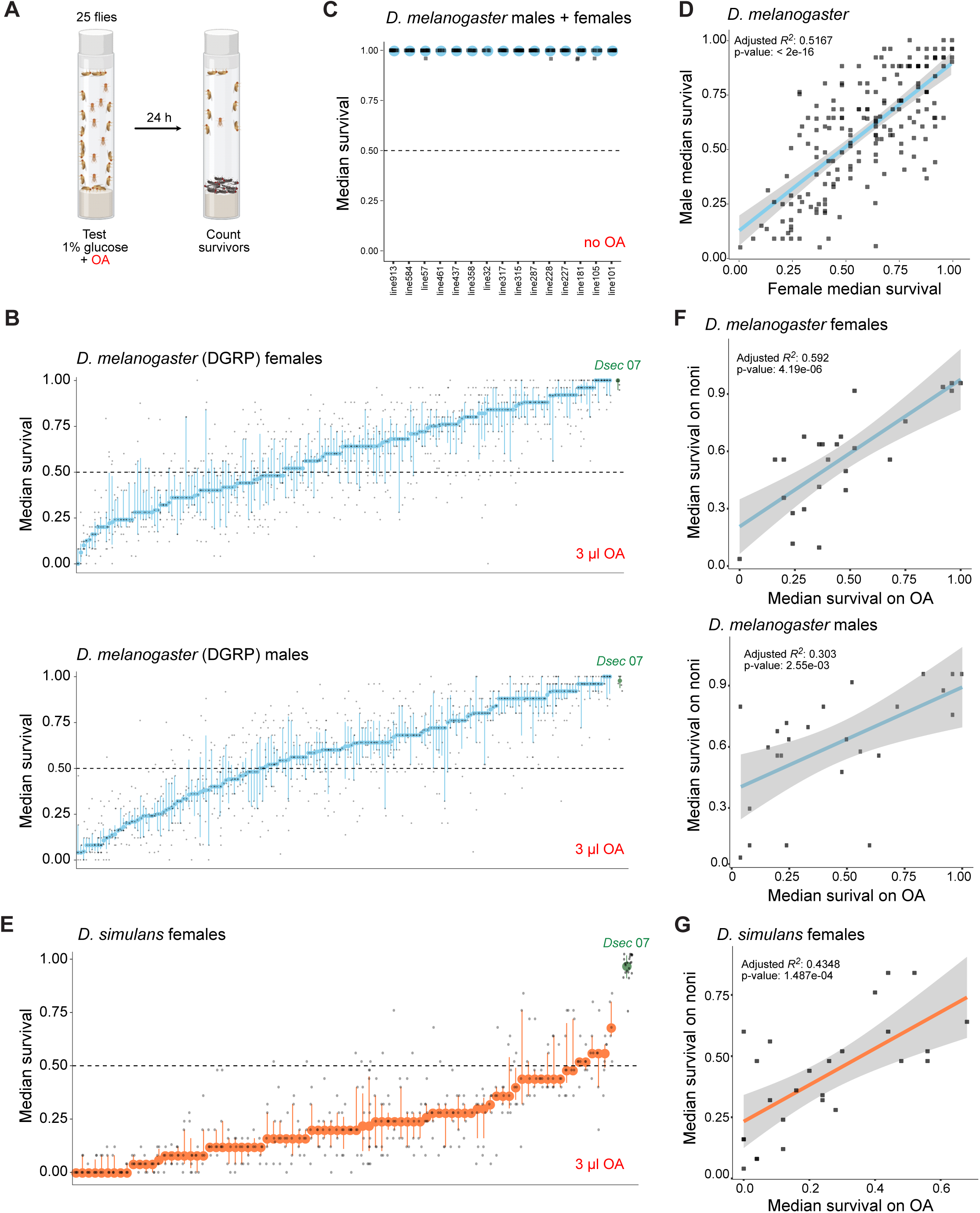
Phenotypic characterization of OA resistance in *D. melanogaster* and *D. simulans* strains. (A) Schematic of the OA resistance tube test. (B) Representation of OA resistance of the *D. melanogaster* (DGRP) strains for females (top) and males (bottom). Strains are arranged on the x-axis by increasing median survival (y-axis). Blue dots are the median values, blue whiskers are the inter-quantile ranges, and grey dots are the individual replicates. The high intrastrain variability is concordant with previous toxicological studies with OA (Amlou et al., 1997; Colson, 2004). Raw data are available in File S1. (C) Median survival of a subset of *D. melanogaster* strains in the absence of OA. (D) Male/female survival correlations based on the median values from (B). Shaded area represents the 95% confidence interval. (E) Representation of the OA resistance of the *D. simulans* strains for females. Strains are arranged on the x-axis by increasing median survival (y-axis). Orange dots are the median values, orange whiskers are the inter-quantile ranges, and grey dots are the individual replicates. Raw data are available in File S1. (F-G) OA/noni fruit survival correlation for a subset of 25 DGRP strains (F) and 25 *D. simulans* strains (G). Shaded area represents the 95% confidence interval. Raw data are available in File S2.

Using this assay, we analyzed resistance to OA in males and females of 186 *D. melanogaster* lines from the DGRP (Table S1). Survival to OA varied remarkably widely across the strains in both sexes, with median survival ranging from nearly 0 to 1, reaching *D. sechellia*-like levels of resistance (at least under these conditions) (Figure 2B). In control assays without OA, using a subset of lines spanning the range of survival probabilities, we found survival of essentially all flies regardless of strain or sex, indicating that the mortality observed under experimental conditions is attributable to OA exposure (Figure 2C). Survival probabilities were strongly correlated between sexes (Figure 2D) and no significant sex differences were revealed in a linear mixed model including sex as a fixed effect and strain as a random effect (β=−0.0128±0.0138, t=−0.924, df=185, p=0.357). To assess a potential effect of age, which might affect the degree of resistance through the maturation state of cuticular hydrocarbons (Kuo et al., 2012), we examined OA resistance of a subset of strains aged between 1-2 days or 3-7 days post eclosion. No difference was observed in any of the tested strains (Figure S1B).

We performed a similar screen of a panel of *D. simulans* strains (Signor et al., 2018) (Table S1). As in *D. melanogaster*, no sex differences were observed in a small subset of lines (Figure S1C), leading us to restrict full screening to females. Across the 84 tested lines, we observed a wide range of survivability, from 0-0.68 median survival probability (Figure 2E). In control assays in the absence of OA, essentially complete survival of all tested lines was observed (Figure S1D).

Together these screens reveal striking phenotypic diversity in OA resistance in both *D. melanogaster* and *D. simulans* lines. The broad distribution of survival probabilities across the panels indicates that susceptibility/resistance can be shaped by numerous genetic factors.

### OA resistance correlates with resistance to noni but not other toxins

To evaluate the ecological relevance of OA resistance observed in these *D. melanogaster* and *D. simulans* panels, we examined 25 *D. melanogaster* and 25 *D. simulans* strains – spanning the observed ranges of OA survivability – for resistance to noni pulp. Survival on noni strongly correlated with survival on OA in both sexes of *D. melanogaster* (Figure 2F), and in *D. simulans* (Figure 2G). These results confirm the central contribution of this chemical to fruit toxicity and, by extension, imply that that genetic determinants of susceptibility/resistance to OA are potentially relevant to underlie susceptibility/resistance to noni.

The DGRP has been extensively characterized for a multitude of phenotypes, including resistance to diverse toxins, such as organophosphates, pyrethroids and neonicotinoids (Battlay et al., 2018; Battlay et al., 2016; Denecke et al., 2017; Duneau et al., 2018; Green et al., 2019; Najarro et al., 2017; Schmidt et al., 2017). We assessed correlations of resistance to OA and these other toxins using DGRPool (Gardeux et al., 2023), but no significant associations were detected. Similarly, in more limited testing of *D. simulans* survival on pyrethrin-based and fatty acid-based insecticides, we did not detect any significant correlation between the degree of resistance of different lines to these chemicals and OA (Figure S1E). These findings imply that the mechanistic basis of OA susceptibility and resistance is distinct from that of other insecticides.

Broader surveying of OA resistance with >500 other traits in DGRPool revealed a weak but significant correlation with alcohol sensitivity (suggestive of partially overlapping toxicity mechanisms) (Table S2); other correlated traits, including male aggression and average lifespan, are harder to explain.

Finally, we tested for correlation between OA resistance and *Wolbachia* infection. *Wolbachia*-infected flies showed slightly higher median survival (estimate = 0.083, p = 0.020), but the effect was very small and explained less than ∼3% of the variation in OA resistance (adj. R² = 0.024). These results are consistent with our axenic fly assays (which should also lack *Wolbachia* infection (Shropshire, 2024)), further supporting the conclusion that the microbiome plays little-to-no role in this phenotype.

### Genome-wide association studies of OA resistance

To investigate the genetic basis of the observed variation in OA resistance, we tested for associations between median resistance to OA (File S1) and genetic variants segregating in the respective species’ panels. For the DGRP, GWAS was performed separately for males and females (Figure S2A) as well as in a combined analysis (Figure 3A). Only one SNP was identified across all three analyses (Figure 3A and Figure S2A); this SNP falls in the coding sequence of the gene *bark beetle* (*bark*, also known as *anakonda*) on chromosome 2L, resulting in an amino acid substitution (I2114V), as discussed further below. The presence of the alternative allele significantly reduced OA resistance in both male and female flies (Figure 3B).

**Figure 3.**
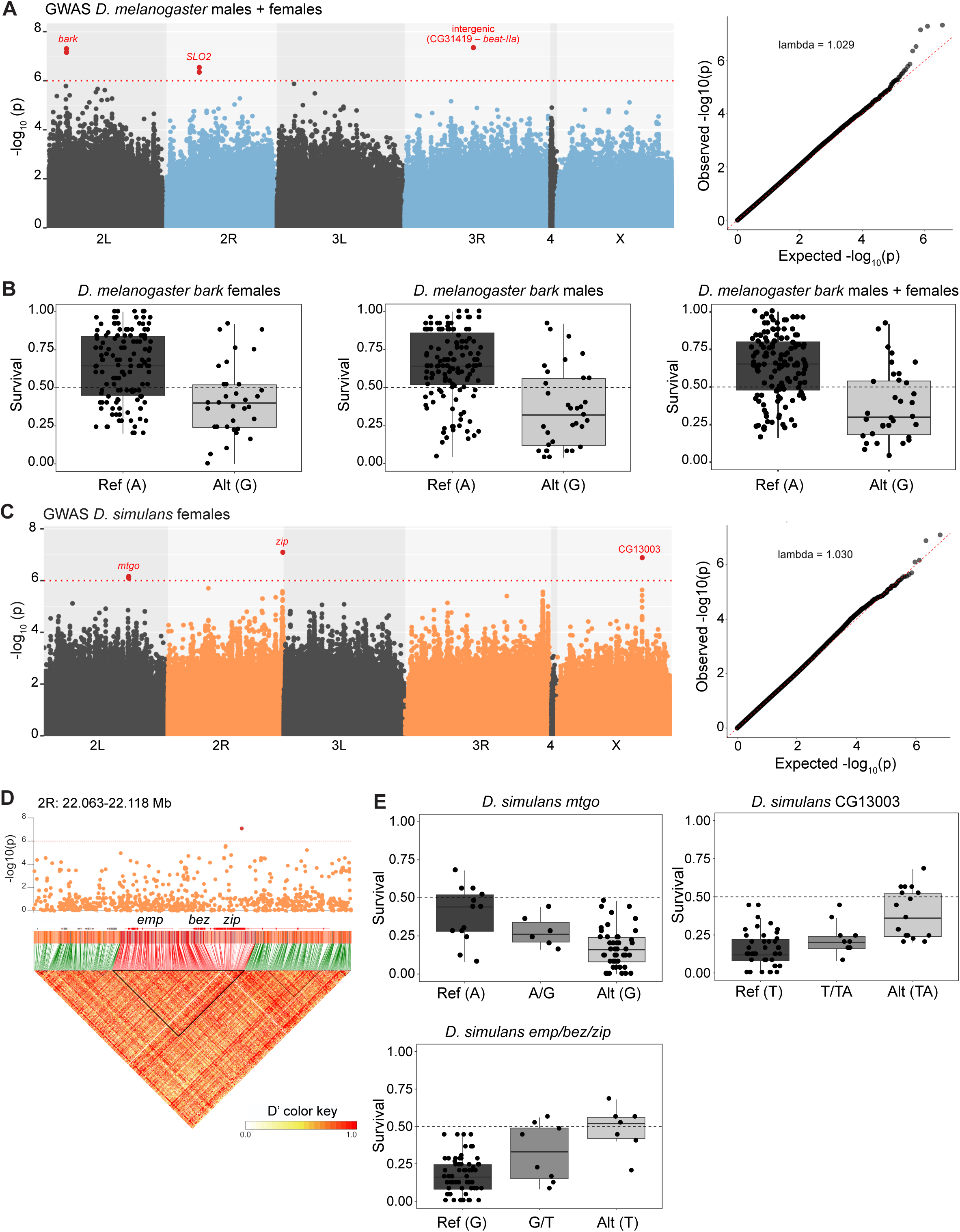
Genome-wide association analysis of OA resistance in *D. melanogaster* and *D. simulans* strains. (A) Left: Manhattan plot of the GWAS of OA resistance, based on median survival of males and females combined. Each dot represents a SNP, plotted on the x-axis according to its genomic position. The y-axis shows -log_10_(p) values for the association between genotype and OA resistance. The red dotted line marks a conservative arbitrary threshold at p ≤ 1×10^-6^. SNPs exceeding this threshold are highlighted in red. Beyond the sole SNP (in *bark*) common to the GWAS of males, females and combined sexes several other significant SNPs were found only in the male and combined analyses and were not considered further in this work. Right: Quantile–Quantile (QQ) plot showing the observed distribution of association test statistics versus the expected distribution under the null hypothesis of no association. Raw data are available in File S3. (B) Boxplots of median OA survival of the tested strains containing the reference (Ref) or alternative (Alt) alleles for the most significant SNP in *bark* for females, males and combined sexes. (C) Same as in (A) but for *D. simulans* females. (D) Zoomed-in Manhattan plot of the 2R region (2R: 22.062 -22.118 Mb) around the significant SNP with underlying linkage disequilibrium (LD) structure. LD is represented as a heatmap (D′ values, ranging from white to red), with genes encompassed within the significant LD block (black triangle) highlighted in red. (E) Boxplots showing median OA survival across tested strains carrying the reference (Ref), heterozygous, or alternative (Alt) alleles for the most significant SNPs identified in (C). The title of each plot indicates the gene(s) encompassed by the corresponding significant SNP.

GWAS of the *D. simulans* panel identified three peaks (Figure 3C). A small peak on 2L falls on the gene *miles to go* (*mtgo*), while the peak on the X chromosome lies in CG13003. The main peak on chromosome 2R is located in an exon of *zipper* (*zip*). However, this locus is part of a 22.5 kb linkage disequilibrium block comprising two additional genes, *epithelial membrane protein* (*emp*) and *between emp and zip* (*bez*) (Figure 3D). Because these genes are in strong linkage disequilibrium, it is not possible to determine whether the association with the phenotype is driven by one specific gene or by multiple genes within this block. For each of these loci, alternative alleles showed an additive effect on OA survivability (Figure 3E). The identified variants all fall in non-coding regions except for the one in *zip,* which causes a synonymous mutation in position R445. The effect of these variants is unclear, and we speculate that their role may be regulatory or that the real causative SNPs were not picked up by our analysis.

### Expression, sequence and functional analysis of candidate OA resistance genes

We proceeded to examine the candidate genes using available information from expression (Fly Cell Atlas (Li et al., 2022), Fly Atlas 2 (Krause et al., 2022)) and functional studies in *D. melanogaster* (from FlyBase (Ozturk-Colak et al., 2024)) (Figure 4A-B and Figure S2B).

**Figure 4.**
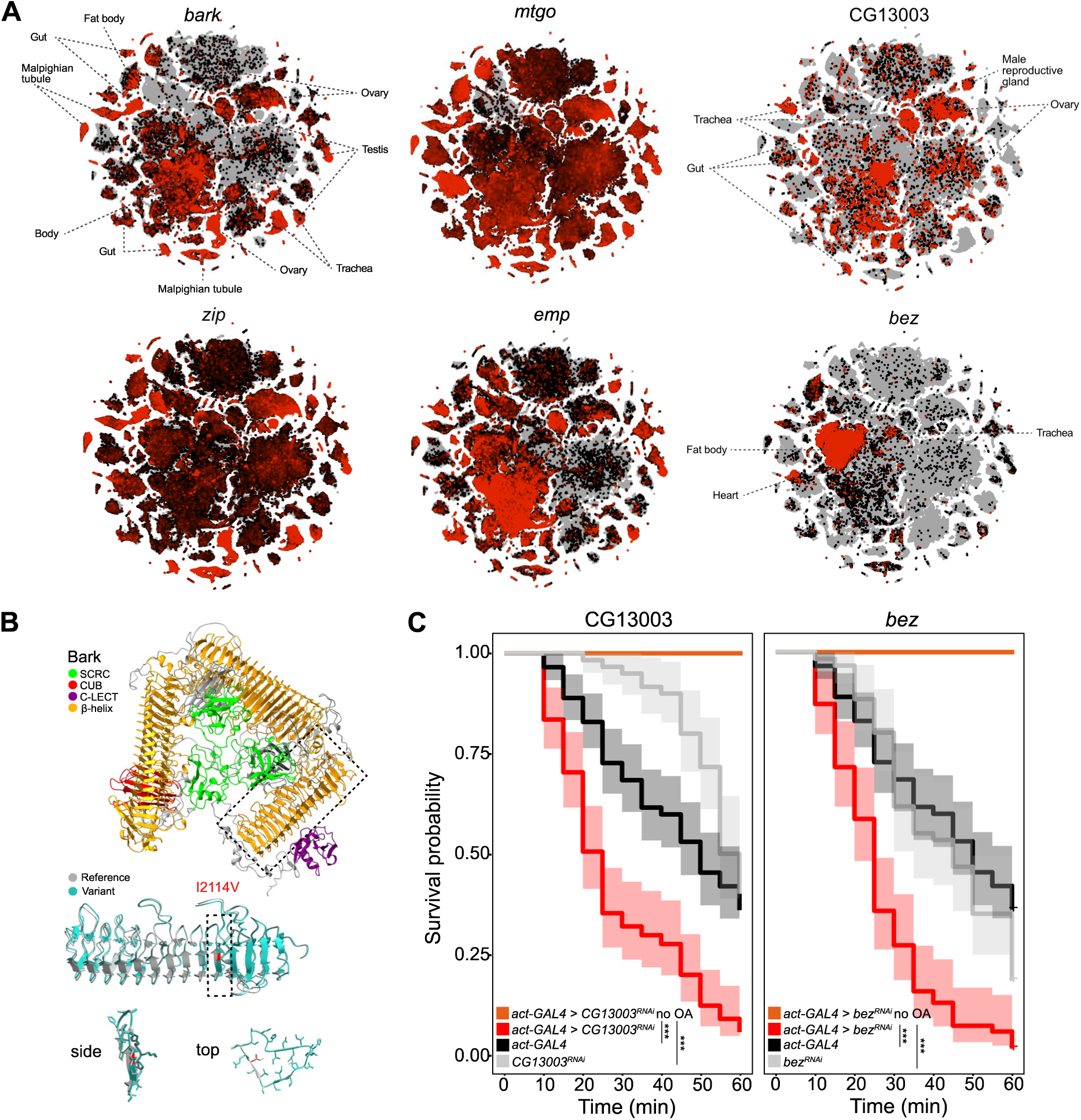
Gene expression, structural modeling, and functional validation of OA resistance genes in *D. melanogaster*. (A) Expression profiles of candidate genes from the Fly Cell Atlas (10x stringent dataset). Relative expression levels are depicted on the default black-to-red scale from SCope interface, with labels marking tissues in which expression is enriched. (B) Top: predicted structure of Bark (based on *D. melanogaster* sequence) carrying the reference variant (I2114). The structure shows the tripartite organization of bark: the β-helix domains form the characteristic β-barrel architecture. Middle: detailed view of the β-helix domain, showing the predicted arrangement of β-helix repeats. The model compares the reference (I2114, grey) and variant (V2114, blue) structures. The location of residue 2114 is marked in red. Bottom: magnified views (side and top) of the β-helix repeat carrying I2114V (red). (C) OA survival curves for and CG13003*^RNAi^* and *bez^RNAi^* flies and the corresponding controls. Each replicate consisted of ten 2-7 day-old female flies. N replicates ≥ 6, n flies ≥ 60. Shaded area represent the 95% confidence interval. Genotypes: *act-Gal4/+;UAS*-*CG13003^RNAi^/+*, *act-Gal4/+*, *UAS-CG13003^RNAi^/+*, *act-Gal4/+;UAS*-*bez^RNAi^/+*, *act-Gal4/+*, *UAS-bez^RNAi^/+*. Raw data are available in File S1.

The sole consistent hit from the *D. melanogaster* GWAS, *bark*, encodes a large (>3000 amino acids) type I transmembrane protein. Based on AlphaFold predictions, the Bark extracellular domain is composed of three repeats of a scavenger receptor cysteine-rich domain and β-helix repeats, and also contains CUB and C-type lectin domains (Figure 4B). *bark* is broadly-expressed in *D. melanogaster*, including the digestive, renal, excretory and reproductive systems (Figure 4A), with the greatest tissue enrichment in the midgut (Figure S2B). Bark has an important function in the maturation of septate junctions between epithelial cells in various tissues (Byri et al., 2015; Hildebrandt et al., 2015). Such junctions have a critical role in cell-cell adhesion and in prevention of diffusion of small molecules through the paracellular space. The variant amino acid position associated with OA resistance (I2114V) is located within one of the β-helix domains of the third repeat. To assess a potential effect of this variant on protein structure, we generated structural models of this domain using both the reference (I2114) and variant (V2114) sequences. Superposition of these models did not reveal a noticeable difference between them (Figure 4B); however, we cannot exclude the possibility that this amino acid variant leads to a subtle conformational change and/or has an effect on protein-protein interactions of Bark.

Of the candidates emerging from the *D. simulans* GWAS, *mtgo* encodes a member of the Fibronectin type III superfamily of cell adhesion proteins (Syed et al., 2019). In *D. melanogaster*, this gene displays near-ubiquitous expression; consistently loss-of-function mutations lead to pupal lethality due, in part, to a developmental requirement at neuromuscular junctions (Syed et al., 2019).

CG13003 is uncharacterized and lacks recognizable protein domains. However it contains an N-terminal signal sequence, suggesting that it is secreted, and is enriched in proline and serine residues (>10% each). Such properties are reminiscent of extracellular matrix proteins, such as mucins (Syed et al., 2008). In *D. melanogaster* larvae, CG13003 is most highly enriched in trachea (Figure S2B), a tissue expression also observed in adults, together with expression in gut epithelia and reproductive tissues (Figure 4A). We speculate that the encoded protein contributes to formation of an extracellular barrier that protects tissues from OA toxicity.

Within the linkage disequilibrium block on chromosome 2R, all three genes (*zip*, *emp* and *bez*) exhibit properties of potential relevance for OA resistance. *zip* is very broadly expressed and encodes a non-muscle myosin heavy chain that is required for regulating the actin cortical network in many cellular and tissue contexts. Within this general role, however, *zip* interacts with septate junction-associated proteins and contributes to acto-myosin contractility that can influence junctional organization (Holland et al., 2025), hinting at the possibility that Zip and Bark affect OA resistance through a related mechanism. *emp* and *bez* both encode members of the CD36 family of transmembrane scavenger receptors, which have diverse ligands and biological roles (Pepino et al., 2014; Silverstein and Febbraio, 2009). *emp* is broadly expressed in epithelial tissues, including trachea, and is an essential gene that is best-characterized for its role in airway morphogenesis (Pinheiro et al., 2023); as for CG13003, this gene might be pertinent for OA resistance should the tracheal system be an entry point for OA into the body. Of particular interest, however, is Bez, which is most prominently expressed in the fat body, where it functions as a fatty acid transporter involved in lipid export from this organ (Carrera et al., 2024). The insect fat body is a key tissue for chemical storage and detoxification (Arrese and Soulages, 2010), raising the possibility that altered function of Bez contributes to OA resistance through enhanced sequestration in this organ.

Manipulation of many of these candidate genes to test their contribution to OA resistance is complicated by their essentiality for animal viability. We therefore focused on CG13003 and *bez*, both because these are the most-selectively expressed of the candidates (Figure 4A), and because these genes were also linked to increased OA resistance in an experimental evolution experiment in *D. simulans* (Marconcini et al., 2025). Transgenic RNAi knock-down of these genes, did not affect survival or lead to obvious morphological or behavioral defects. However, when tested for resistance to OA both CG13003*^RNAi^* and *bez^RNAi^* animals displayed a significant reduction in survivability compared to control genotypes (Figure 4C). These results suggest that the encoded proteins contribute to conferring resistance, rather than susceptibility, to this toxin. Although these data support the promise of the GWAS in identifying functionally relevant genes, future studies will be required to assess whether these and other genes are also relevant for OA resistance in *D. sechellia* and to elucidate their precise mechanism of action.

### Transcriptome-to-phenotype association identifies additional candidate OA resistance genes

While GWAS is powerful for pinpointing SNPs associated with a trait, a transcriptome-to-phenotype study allows the identification of differences in gene expression that might be functionally relevant. The availability of whole-fly, bulk RNA-sequencing datasets for the DGRP (Huang et al., 2015), has enabled transcriptome-to-phenotype analyses on this panel for various insecticide resistance traits, identifying several detoxification-related genes (Battlay et al., 2016; Denecke et al., 2017; Green et al., 2019). Profiting from this dataset, we fitted a linear model regressing median resistance to OA to the mean expression of 18,140 transcripts. This analysis yielded 52, 16, and 75 candidate transcripts for males, females, and the combined dataset, respectively, that display an association between expression level and OA resistance (Figure 5A). We could not find common hits with our GWAS analyses. Gene ontology analysis using PANGEA (Hu et al., 2023) revealed enrichment for a range of terms, including metabolic processes (e.g., the NADH dehydrogenase ND-75 and the cytochrome P450 Cyt-c1), mitochondrial function (e.g., ATP synthase subunit D and Superoxide dismutase 2) and cytoskeleton organization (e.g., Act88F and Tm2) (Figure 5B). The enrichment of these pathways matches well with those identified in complementary genetic analyses of OA resistance through experimental evolution and CRISPR-based loss-of-function screens in cell lines (Marconcini et al., 2025).

**Figure 5.**
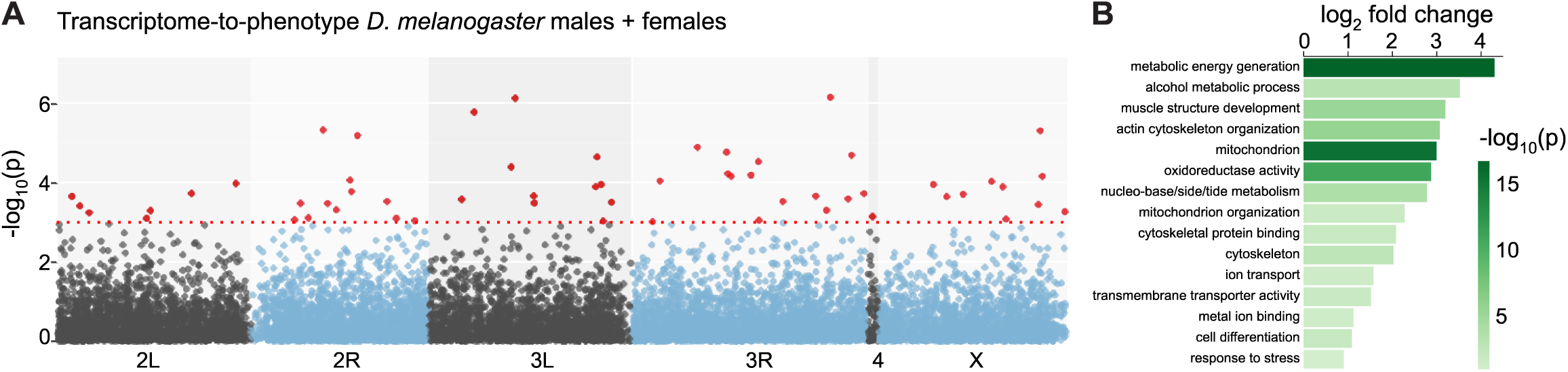
Transcriptome-to-phenotype associations for OA resistance in DGRP strains. (A) Manhattan plot of the transcriptome-to-phenotype association analysis in the DGRP, based on median expression values combining males and females. Each gene is represented as a single dot positioned along the x-axis according to its genomic location. The y-axis shows the -log_10_(p) for the association between expression and OA resistance. The red dotted line marks an arbitrary significance threshold of p ≤ 1×10^-3^. Transcripts exceeding this threshold are highlighted in red. (B) PANGEA gene ontology analysis of the significant genes identified in (A) using the curated Drosophila GO Subsets. Raw data are available in File S4.

## Discussion

The evolution of insect resistance to toxins is often linked most strongly with mutations in genes encoding target proteins, with striking examples of parallel or convergent evolution between different insect species exposed to natural or artificial toxins that have the same target (Busvine, 1951; Dong, 2007; Guo et al., 2023). Resistance to the toxin OA appears to be fundamentally different. This trait was historically first-recognized in insects because of the striking, high resistance exhibited by *D. sechellia*, which is a critical adaptation to its specialized ecological niche (Amlou et al., 1997; Auer et al., 2021; Legal et al., 1994; R’Kha et al., 1991). Mapping studies reveal a complex genetic architecture of *D. sechellia*’s OA resistance (see Introduction). More recently, a strain of *Drosophila yakuba* – a more distant relative within the *melanogaster* subgroup of drosophilids – was found to exhibit similar ecological specialization on noni fruit and displays higher OA resistance than the parental species (Yassin et al., 2016). Using a genome-wide scan of targets of natural selection across *D. yakuba* populations, several loci associated with this recent specialization on noni were identified, but these displayed only limited overlap with genomic regions identified in *D. sechellia*. In this work, we show that even different inbred strains of *D. melanogaster* and *D. simulans* exhibit large variation in OA susceptibility/resistance, and capitalize on such phenotypic variation to identify, through association studies, several novel candidate genes contributing to this trait.

Integrating the genes identified in this work with those that emerged from previous studies of OA resistance in various drosophilid species (Table 1 and Introduction), reinforces the evidence for the contribution of several pathways to this trait. The nature of the encoded proteins and expression patterns points to a complex interplay of multiple tissues and molecular processes underlying resistance to OA. Such breadth suggests that OA does not have a singular cellular or molecular target, but might affect a basic cellular process, an idea supported by the toxicity of OA to cultured insect cell lines (Kaczmarek et al., 2024; Kaczmarek et al., 2022; Marconcini et al., 2025), yeast (Mota et al., 2024) and bacteria (Liu et al., 2013). The candidate resistance genes seem less likely to be a target of OA but rather contribute to a variety of defense mechanisms, including cuticular and epithelial barrier functions, gut/renal system chemical detoxification, lipid transport, and general metabolic activity. Identification of Bark from our *D. melanogaster* GWAS is of note, as it highlights a novel defense mechanism of insects against toxic chemicals. Oral insecticides are thought to have to cross the midgut epithelium to be able to exert their toxic effects (Denecke et al., 2018), but there has been almost no previous consideration for how natural variation in the integrity of this internal tissue barrier – analogous to the external cuticular barrier – might determine susceptibility or resistance to toxins (Chen et al., 2023). Examination of the contributions of other genes involved in septate junction formation to resistance to OA, as well as other toxic chemicals, will be of interest.

**Table 1.** Summary of properties of candidate genes underlying OA resistance.

| Gene | Protein domains | Putative/known cellular role(s) | Notable tissue expression | Species | Stage | Methods / Supporting data | References |
| --- | --- | --- | --- | --- | --- | --- | --- |
| *Alkbh7* | α-ketoglutarate-dependent dioxygenase | mitochondrial RNA demethylation; programmed necrotic cell death; fatty acid metabolism | ubiquitous | *Dmel/Dsim/Dsec* | adult | Experimental Evolution, CRISPR, RNAi, KO, Expression analyses | (Marconcini et al., 2025) |
| *bark* | SCRC, CUB, C-LECT, β-helix | septate junction maturation | broad | *Dmel* | adult | GWAS | *This study* |
| *bez* | CD36 | fatty acid transport | fat body | *Dsim/Dmel* | adult | GWAS, RNAi | (Marconcini et al., 2025), *this study* |
| **CG13003** | Mucin-like? | unknown | trachea | *Dsim/Dmel* | adult | GWAS, RNAi | (Marconcini et al., 2025), *this study* |
| *emp* | CD36 | fatty acid transport | ubiquitous | *Dsim* | adult | GWAS | *This study* |
| *Est-6* | Type-B carboxylesterase /lipase | metabolic detoxification | broad | *Dmel* | adult | RNA-seq and RNAi | (Lanno et al., 2019a) |
| *gce* | Myc-type, bHLH PAS | juvenile hormone receptor | ubiquitous | *DsecxDsim* | larva | QTL | (Huang and Erezylmaz, 2015) |
|  |  |  |  | *Dyak* | adult | Population Branch Excess | (Yassin et al., 2016) |
| **GST-E genes** | Glutathione S-transferase | metabolic detoxification | broad | *Dsec* | adult | Microarray | (Dworkin and Jones, 2009) |
|  |  |  |  | *Dyak* | adult | Population Branch Excess | (Yassin et al., 2016) |
| *kraken* | Serine hydrolase | metabolic detoxification | excretory system | *Dmel/Dsim/Dsec* | adult | Experimental Evolution, CRISPR, RNAi, KO, Expression analyses | (Marconcini et al., 2025) |
| *mtgo* | Fibronectin type III | neuromuscular junction branching and growth | ubiquitous | *Dsim* | adult | GWAS | *This study* |
| *Osi6, Osi7, Osi8* | DUF1676 | endosomal-localized; regulation of cuticle secretion/shaping | epithelia, trachea | *Dyak* | adult | Introgression with CAPS markers | (Hungate et al., 2013) |
|  |  |  |  | *DsecxDsim* | adult | Population Branch Excess | (Yassin et al., 2016) |
|  |  |  |  | *Dsec* | larva, adult | RNA-seq | (Lanno et al., 2017; Lanno et al., 2019b) |
|  |  |  |  | *Dmel* | larva, adult | RNAi | (Andrade Lopez et al., 2017; Lanno et al., 2019b) |
| **serine protease genes** | Peptidase S1 family serine proteases | detoxification | reproductive organs, epithelia | *Dyak* | adult | Population Branch Excess | (Yassin et al., 2016) |
| **Tweedle genes** | DUF243 | chitin-binding, cuticle development | epithelial, trachea | *DsecxDsim* | larva | QTL | (Huang and Erezylmaz, 2015) |
|  |  |  |  | *Dyak* | adult | Population Branch Excess | (Yassin et al., 2016) |
| *zip* | Non-muscle myosin 2 heavy chain | cortico-actin dynamics | ubiquitous | *Dsim* | adult | GWAS | *This study* |

The limited overlap in the specific genes identified between individual studies (Table 1) reinforces the idea of no singular major-effect gene contributing to OA resistance. One explanation for the lack of common hits both between the present *D. melanogaster* and *D. simulans* GWAS, and with previous investigations, is that GWAS can only recover associations from the finite variation segregating in a given panel of strains. Thus, different starting populations will yield different sets of candidate genes. We also note that there is also only partial overlap between the *D. simulans* GWAS with genetic variants identified in our previous experimental evolution of *D. simulans* populations (Marconcini et al., 2025), despite the base populations being initially generated from the same lines profiled here. The failure to identify, for example, *kraken* in the GWAS might be because of insufficient variability of this gene across the lines, a small effect size, or a detectable contribution of this gene only in a particular genetic background (of both strain and species). More generally, the contributions of different genes to OA susceptibility/resistance is also likely to depend upon the life stage, the dose of this chemical and the method of exposure, one or more of which factors vary between different studies.

The limited overlap of specific genes between studies in different strains/species also suggests there are many possible paths towards OA resistance. It is nevertheless notable that standing genetic variation in populations of *D. melanogaster* and *D. simulans* – both isolated from restricted geographic localities – is sufficient to encode a broad phenotypic range in resistance to OA. This observation hints at the possibility that OA resistance in *D. sechellia* also emerged, at least initially, from genetic variants already present in the ancestor of *D. sechellia* and *D. simulans* (and perhaps also *D. melanogaster*) rather than through *de novo* mutations of large-effect size.

Given the evident genetic complexity of this superficially simple trait, it is clearly useful to combine the results of distinct approaches – each with their advantages and disadvantages – to identify as many candidate genes as possible underlying this phenotype. Further work will be required to determine whether genetic associations reflect causal requirements, both in laboratory inbred strains and in strains in nature, and the precise site and mechanism of action of the encoded proteins.

## Methods

### Drosophilid strains and culture

All drosophilid strains used in this study were reared on wheat flour/yeast/fruit juice medium at 25°C under a 12 h:12 h light:dark cycle. Wild-type and transgenic strains are listed in Table S1.

Axenic stocks were generated following standard protocols (Charroux and Royet, 2020). In brief, we maintained stocks for 3-5 generations on standard medium supplemented with an antibiotic cocktail (50 µg/ml kanamycin (Applichem, CAS 5965-95-7), 50 µg/ml ampicillin (Applichem, CAS 69-52-3), 10 µg/ml tetracycline (Sigma Aldrich, CAS 64-75-5), and 5 µg/ml erythromycin (Carl Roth, CAS 114-07-8)) until germ-free. To assess the absence of fly-associated microbes, ten flies were homogenized in 600 µl sterile PBS. From each homogenate, 100 µl were plated in triplicate on different microbiological media: Plate Count Agar (30°C), MRS agar (37°C), D-mannitol agar (30°C), and LB agar (37°C). Plates were incubated for 48 h to detect potential bacterial growth.

### OA resistance assay

OA resistance was assessed in plates or tubes essentially as described (Marconcini et al., 2025). In brief, for the plate assay, different volumes of OA were applied to the inside of the lid of a Petri dish (60 mm diameter, 15 mm height; Greiner Bio-One). 10 female flies were transferred to each plate after brief CO_2_ anesthesia for sorting. Flies were allowed to recover (typically 5 min), and mortality was then recorded every 5 min for 60 min. Statistical analysis of the survival curves was conducted fitting a mixed effect Cox regression model as implemented in the R package coxme (Therneau, 2024). Genotype was included as a fixed effect, and replicate dishes were modeled as a random effect. For axenic flies, the volume of OA was also included as a fixed effect.

For the tube assay, absorbent paper (Kimtech, 7552) was placed at the bottom of empty vials (25 mm diameter, 95 mm height, Milian SA), to which 1 ml of 3% D-glucose (CAS 14431-43-7, neoFroxx) solution was added, to prevent starvation and desiccation. 3 µl of OA (CAS 124-07-2; Sigma; >99% purity) was applied to the paper, and the vials were immediately sealed with cotton caps to minimize vapor loss. Flies (aged 1-7 days) were anesthetized with CO_2_, sexed, and transferred in groups of 25 males or 25 females per vial. Vials were placed upside down for 5 min to allow recovery, then maintained for 24 h at 25°C under a 12 h:12 h light:dark cycle. Alive flies were counted at the end of the assay, and survival probability was calculated as the number of survivors/total number of flies. Control assays followed the same procedure, without the addition of OA. To test for sex differences in OA survival, we fitted a linear mixed-effects model with sex as a fixed effect and line as a random effect. Covariation between sexes was also examined by assessing the correlation between male and female median survival across lines.

For a subset of *D. simulans* strains, the tube assay was also used to assess resistance to two commercially-available insecticides: the pyrethrin-based “Spruzit” AF (Neudorff, W-6670) (700 µl) and the fatty acid-based “Natural” (Andermatt Biogarten, W-6107) (100 µl). The volumes of these insecticides were chosen following pilot trials with a few *D. simulans* strains, representing a dose that leads to a range of survival probabilities.

### Noni fruit resistance assay

Noni fruit was harvested from *M. citrifolia* plants cultivated in the University of Lausanne greenhouses. Noni pulp purée was prepared by homogenizing five ripe fruits and stored at -20°C until use. Resistance was assessed in the tube assay as described above, replacing OA and the tissue paper soaked with glucose with 2 g of thawed noni pulp purée. A previous study indicated that noni contains 3.06 g OA/kg fruit (Pino et al., 2009); assuming an equivalent quantity in our fruits, 2 g noni pulp should contain 6.73 µl OA, approximately two-fold higher than the amount of pure OA used. For the tested *D. melanogaster* and *D. simulans* lines, the correlation between median OA resistance and median noni resistance across lines was assessed using a linear model fitted with the lm() function in R 4.0.3 (R Core Team, 2021).

### GWAS

*D. melanogaster:* for the DGRP, we downloaded the binary files from the DGRP2 website (http://dgrp2.gnets.ncsu.edu/data.html). The median survivability scores to OA were used to test for genotype-phenotype association. We fitted a linear model in Plink2.0 using the following parameters: --glm hide-covar --quantile-normalize --variance-standardize --geno 0.2 --maf 0.05. We controlled for population structure adding the --covar parameter using the first 20 PCs obtained with --pca 20. We also controlled for known inversions (i.e., *In(2L)t*, *In(2R)NS*, *In(2R)Y1*, *In(2R)Y2*, *In(2R)Y3*, *In(2R)Y4*, *In(2R)Y5*, *In(2R)Y6*, *In(2R)Y7*, *In(3L)P*, *In(3L)M*, *In(3L)Y*, *In(3R)P*, *In(3R)K*, *In(3R)Mo*, *In(3R)C*) and *Wolbachia* infection modelling them as covariates. The analysis was conducted both on the combined dataset of males and females, as well as separately for each sex.

*D. simulans:* the raw genomic data were downloaded from SRA (SRP075682), aligned to the *D. simulans* reference genome (NCBI GCF_016746395.2) using bwa-mem (Li, 2013) and the results were sorted and compressed using samtools (Danecek et al., 2021). Duplicate reads were then removed with Picard MarkDuplicates (https://broadinstitute.github.io/picard/). Variants were identified using GATK Haplotypecaller (McKenna et al., 2010). We filtered variants with quality lower than 30 with vcftools 0.1.17 (Danecek et al., 2021). Binary input files for the GWAS analysis were generated with plink2.0 (Purcell, 2020; Purcell et al., 2007). The median survivability scores to OA were used to test for genotype-phenotype association. We fitted a linear model in Plink2.0 following the same procedure as for the DGRP. Linkage disequilibrium maps of focal regions were generated with LDBlockShow using the Solid Spine method based on D’ statistics (Dong et al., 2021).

### Transcriptome-to-phenotype association

Whole-fly, bulk RNA-seq data for DGRP lines (Huang et al., 2015) were downloaded from the DGRP2 website (http://dgrp2.gnets.ncsu.edu/data.html), and mean expression across replicates was computed for each transcript. We fitted a linear model between the median survivability scores to OA and each transcript, as in (Battlay et al., 2018; Green et al., 2019), for the 161 lines for which both phenotypic and transcriptomic data are available. A p-value significance threshold of 1×10^-3^ was applied, following (Battlay et al., 2018). Analyses were conducted both on the combined dataset of males and females, as well as separately for each sex. The results were visualized using a Manhattan plot, with each analyzed transcript mapped to the genomic position of its corresponding gene.

### Protein structure prediction

Three-dimensional structure predictions of Bark variants (i.e., with I2114 or V2114, within the longest-encoded protein isoform) were generated using the implementation of AlphaFold2 (Jumper et al., 2021) on the ColabFold platform (https://colab.research.google.com/) (Mirdita et al., 2022), using default parameters. Five ranked models were generated for each sequence, and the top-ranked structure, based on the predicted Local Distance Difference Test (pLDDT) score, was selected for further analysis. The variant protein structures were aligned and visualized in ChimeraX.

## Acknowledgements

We acknowledge the Bloomington *Drosophila* Stock Center (NIH P40OD018537) for *D. melanogaster* stocks. We thank to Blaise Tissot-Dit-Sanfin for cultivation of the *M. citrifolia* plants, and Steeve Cruchet for maintenance of *D. simulans* strains. We thank members of the Benton laboratory for comments on the manuscript. A.M. is supported by an EMBO Long-term Fellowship. Research in R.B.’s laboratory is supported by the University of Lausanne, an ERC Advanced Grant (833548) and the Swiss National Science Foundation (310030_219185). Figure cartoons were created with Biorender.com and SciDraw (10.5281/zenodo.4421109).

## Author contributions

M.M. conceived the project and performed all experiments and analyses in the paper except for the *D. melanogaster* GWAS, which was performed by C.F, who also performed pilot experiments with axenic flies. A.M. generated the axenic fly lines. R.B. conceived and supervised the project. M.M. and R.B. wrote the manuscript, with contributions from C.F. All authors read and approved the final manuscript.

## Supplementary Information

**Table S1. *Drosophila* strains**

**Table S2. Top 10 phenotypic correlations with OA resistance in the DGRP.**

**Figure S1.**
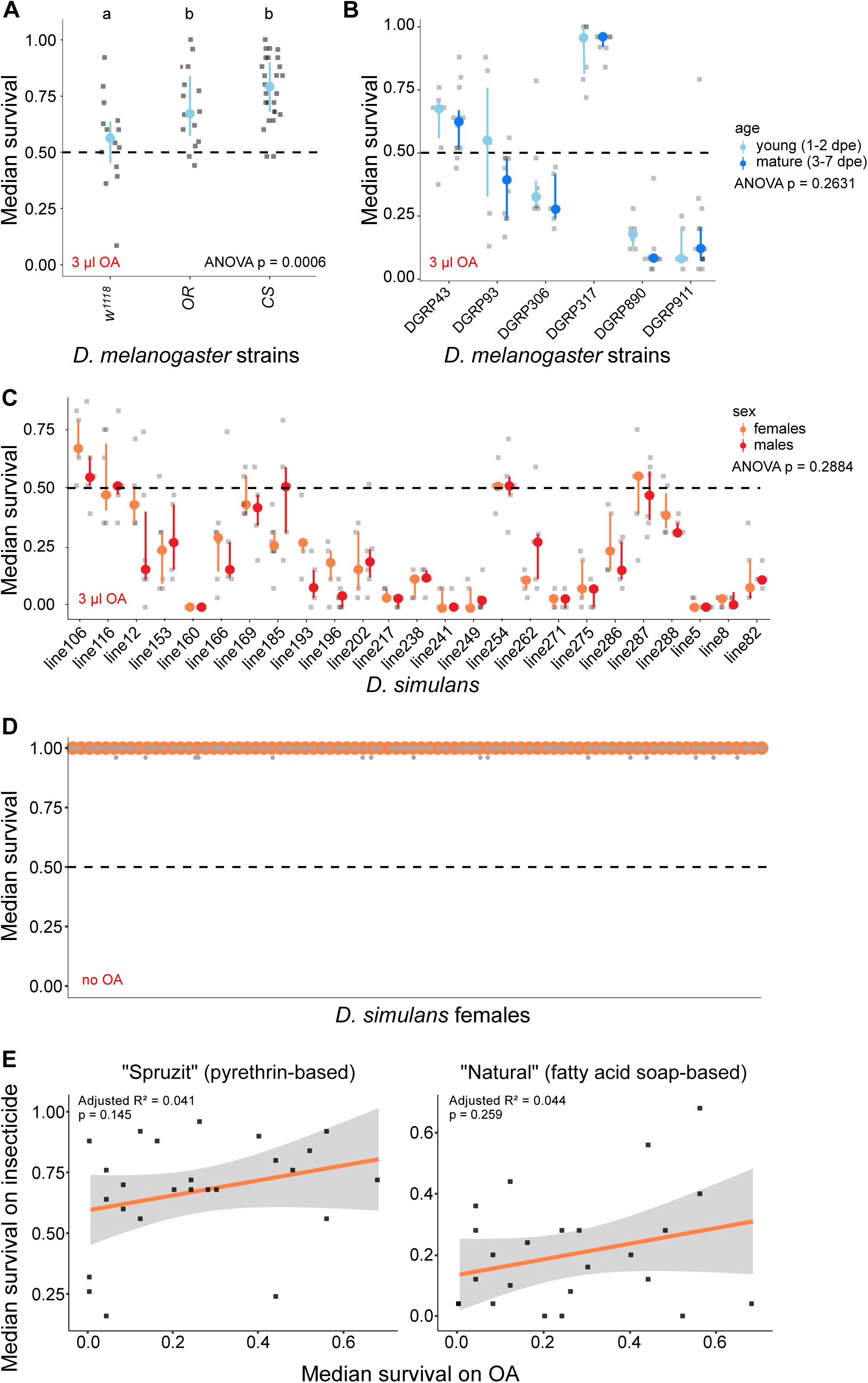
Additional phenotypic characterization of OA resistance in *D. melanogaster* and *D. simulans*. (A) OA resistance of common *D. melanogaster* laboratory strains (*w^1118^*, Oregon-R, and Canton-S) for males and females combined. (B) OA resistance of a subset of DGRP strains in young (1-2 days post-eclosion, dpe) and mature (3-7 dpe) mixed sex flies. (C) OA resistance of males and females of a subset of *D. simulans* strains. (D) Median survival of *D. simulans* strains in the absence of OA. For (A-D) Colored dots represent median values, colored whiskers indicate interquartile ranges, and grey dots correspond to individual replicates. Statistical differences were assessed using ANOVA followed by Tukey’s post hoc test. Raw data are available in File S1. (E) OA/insecticide (containing either pyrethrin or a mixture of medium chain fatty acids) median survival correlations for a subset of *D. simulans* strains. Raw data are available in File S2.

**Figure S2.**
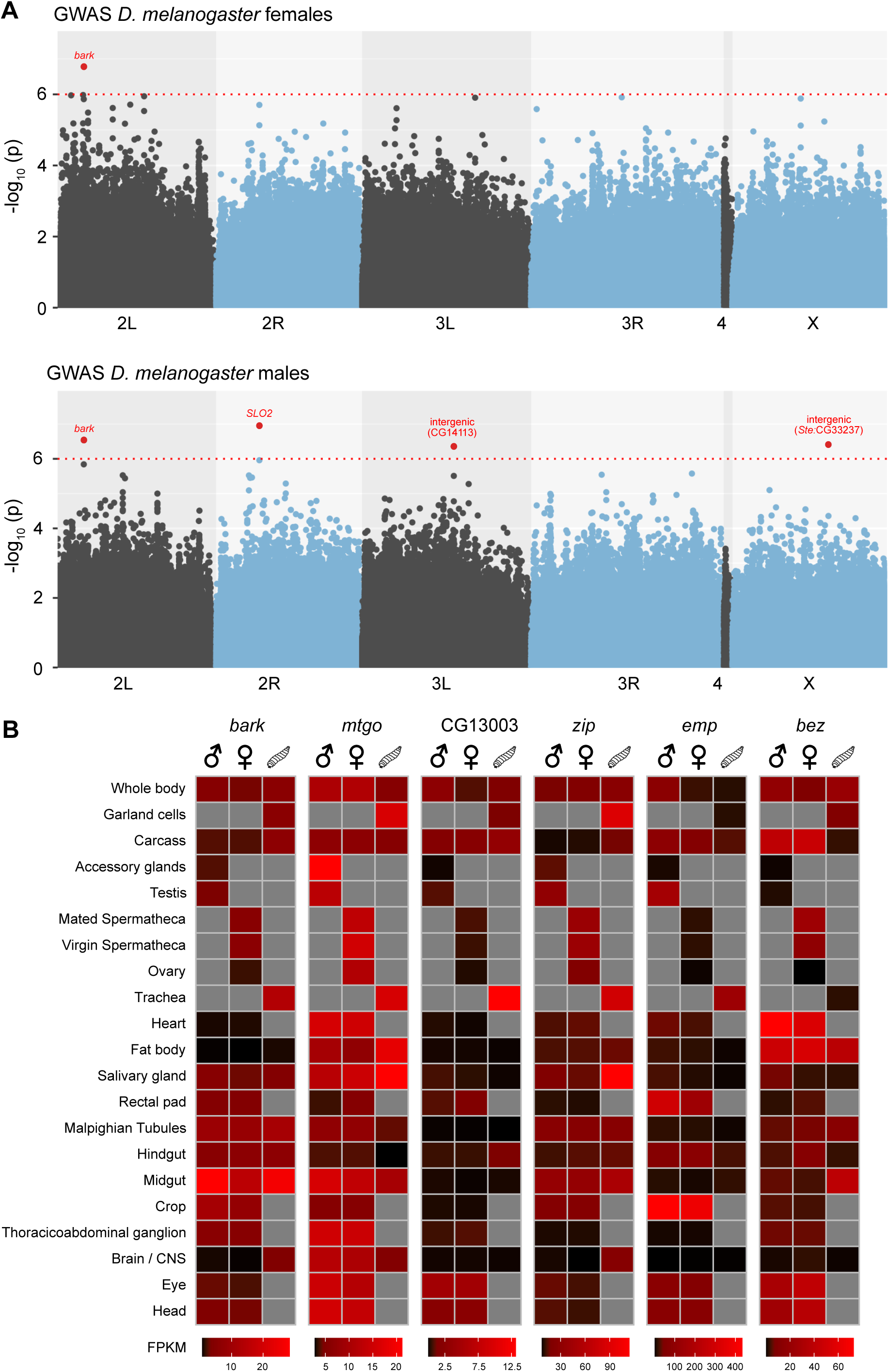
Genetic and expression profiles associated with OA resistance in male and female *D. melanogaster*. (A) Manhattan plot of the GWAS of OA resistance in *D. melanogaster* (DGRP), based on median survival of females (top) and males (bottom). Each dot represents a SNP, plotted on the x-axis according to its genomic position. The y-axis shows -log_10_(p) values for the association between genotype and OA resistance. The red dotted line marks a conservative arbitrary threshold at p ≤ 1×10^-6^. SNPs exceeding this threshold are highlighted in red. Raw data are available in File S3. (B) Heatmap showing expression profiles of the candidate genes across tissues, based on FlyAtlas 2 data. Relative expression levels are depicted on a black-to-red scale. Grey indicates no data are available.

**File S1. OA resistance data for *D. melanogaster, D. simulans* and *D. sechellia*.**

**File S2. OA, noni and insecticide survival probability correlation data.**

**File S3. GWAS results.** *available at*: https://drive.google.com/file/d/1_AoLrDwEnmwIfFyR48JnvXIxdCq2QNzw/view?usp=drive_link

**File S4. Transcriptome-to-phenotype association results and PANGEA analysis.**

